# Dynamics and turnover of memory CD8 T cell responses following yellow fever vaccination

**DOI:** 10.1101/2021.01.23.427919

**Authors:** Veronika I. Zarnitsyna, Rama S. Akondy, Hasan Ahmed, Don J. McGuire, Vladimir G. Zarnitsyn, Mia Moore, Philip L. F. Johnson, Rafi Ahmed, Kelvin Li, Marc Hellerstein, Rustom Antia

**Author notes:** These authors contributed equally to this work (Veronika I. Zarnitsyna, Rama S. Akondy, Hasan Ahmed). & (Veronika I. Zarnitsyna, Rustom Antia).

## Abstract

Understanding how immunological memory lasts a lifetime requires quantifying changes in the number of memory cells as well as how their division and death rates change over time. We address these questions by using a statistically powerful mixed-effects differential equations framework to analyze data from two human studies that follow CD8 T cell responses to the yellow fever vaccine (YFV-17D). Models were first fit to the frequency and division rates of YFV-specific memory CD8 T cells 42 days to 1 year post-vaccination. A different dataset, on the loss of YFV-specific CD8 T cells over three decades, was used to assess out of sample predictions of our models. The commonly used exponential and bi-exponential decline models performed relatively poorly. Models with the cell loss following a power law (exactly or approximately) were most predictive. Notably, using only the first year of data, these models accurately predicted T cell frequencies up to 30 years post-vaccination. Our analyses suggest that division rates of these cells drop and plateau at a low level (0.001 per day, ~double estimates for naive T cells) within one year following vaccination, whereas death rates continue to decline for much longer. Our results show that power laws can be predictive for T cell memory, a finding that may be useful for vaccine evaluation and epidemiological modeling. Moreover, since power laws asymptotically decline more slowly than any exponential decline, our results help explain the longevity of immune memory phenomenologically.

**Author summary:** Immunological memory, generated in response to infection or vaccination, may provide complete or partial protection from antigenically similar infections for the lifetime. Memory CD8 T cells are important players in protection from secondary viral infections but quantitative understanding of their dynamics in humans is limited. We analyze data from two studies where immunization with the yellow fever virus vaccine (YFV-17D) generates a mild acute infection and long-term memory. We find that: (i) the division rate of YFV-17D-specific CD8 T cells drops and stabilizes at ~ 0.1% per day during the first year following vaccination whereas the death rate declines more gradually, and (ii) the number of these cells declines approximately in accordance with a power law (∝time^−0.82^) for at least several decades following vaccination.

## Introduction

Immunological memory is a central feature of the adaptive immune response that underlies successful vaccination, with memory CD8 T cells aiding in faster clearance of subsequent infections. How do division and death rates of expanded pathogen-specific CD8 T cells change with time after pathogen is cleared? Can we predict the number of pathogen-specific cells decades after infection or vaccination based on the early immune response in humans? These are the two main questions that we aim to address here.

Studies in mice following infections with viruses and bacteria, such as LCMV and Listeria monocytogenes, have greatly contributed to our understanding of immunological memory [1–9]. Infections typically stimulate a rapid burst of proliferation of virus-specific CD8 T cells and the generation of a large population of effector cells that clear the virus. Subsequently, more than 90% of the virus-specific pool of CD8 T cells dies, and by day 30 post-infection there is a stable population of long-term memory cells [3–5,8]. Turnover studies, first with BrdU [1, 2] and subsequently with CFSE [6] showed that the memory cell population is maintained by the continuous turnover of cells with stochastic division (with an average time of about 50 days in mice) being balanced by death [6, 7, 9].

Extending studies of immunological memory from mice with relatively short lifespans of 2-3 years to humans (lifespans 70-100 years) has been particularly challenging. It has been shown that the overall number of virus-specific memory CD8 T cells after vaccination slowly declines through human life with an estimated half-life of T cell immunity of 8-15 years [10, 11], and functional memory T cells can be detected for several decades after acute viral infection or immunization with live attenuated virus vaccines [12, 13]. A number of studies have analyzed the turnover of bulk CD8 T cells in humans (typically based on deuterium incorporation) using different cell surface markers which have been associated with populations of naive, effector, memory and stem cell-like memory phenotypes [14–21]. A potential problem with this approach is the inability to unambiguously sort cells at different stages of T cells differentiation by using a few cell surface markers [22–24]. An alternative approach, and the one that we use in this paper, is to focus on a defined population of antigen-specific cells following immunization of individuals naive to a virus in the vaccine. We consider the merits and problems with these different approaches in the Discussion.

In this study, we analyze the dynamics and turnover of CD8 T cells following immunization of individuals with the live attenuated yellow fever vaccine (YFV-17D) [25], which generates a mild acute infection and confers long-term immunity. Our analysis integrates data from two studies. The study by Akondy et al. [23] (hereafter referred to as the Akondy study) *longitudinally* follows the number and turnover of YFV-specific CD8 T cells in a group of 9 individuals during the first year after immunization with YFV-17D vaccine. The number of CD8 T cells specific for an HLA-A2 restricted immunodominant epitope in the NS4B protein (A2-NS4B^214^) of the virus was determined using corresponding peptide-MHC class I tetramers (pMHCs) [22, 23]. The turnover of these cells was measured through labelling of their DNA following administration of heavy water (D_2_O) with subsequent analysis of the die-away kinetics of deuterium in the DNA of these cells after D_2_O administration was stopped. The study by Fuertes-Marraco et al. [26] (hereafter referred to as the Marraco study) was a *cross-sectional* survey that measured the number of YFV-specific T cells using A2-NS4B^214^ tetramers over a much longer time period ranging from few months to three decades following YFV-17D vaccination.

We first build on the Akondy study for the dynamics and turnover of YFV-specific CD8 T cells during the early memory phase of the response. In particular, we quantify how the turnover rate of YFV-specific CD8 T cell population declines over time, and show that this decline is consistent with several models for changes in division and death rates of these cells. We then use the data from the Marraco study for model discrimination. Finally, we discuss how we can predict the number of virus-specific cells decades after vaccination using the power law and the data from the first year T cell responses.

## Results

### Dynamics of the YFV-response during the first year

Immunization with YFV results in a mild acute infection with a peak of virus replication at around 5 to 7 days and total duration about 9 to 14 days (Fig 1A plots data from [23], also see [27] for details of the virus dynamics in different individuals). YFV-specific cells, measured using A2-NS4B^214^ tetramers, proliferate rapidly during the first 21-28 days post immunization, and subsequently their numbers gradually decline over time. The turnover of YFV-specific cells was determined using deuterium labelling. Individuals were given heavy water for the first two weeks following vaccination, which resulted in labelling of the DNA of dividing cells. After administration of deuterium labelled water was stopped, there was a rapid loss of D_2_O in serum (half-time approximately 9 days [23]). The changes in the amount of deuterium in the DNA of YFV-specific cells are shown in Fig 1B.

**Fig 1.**
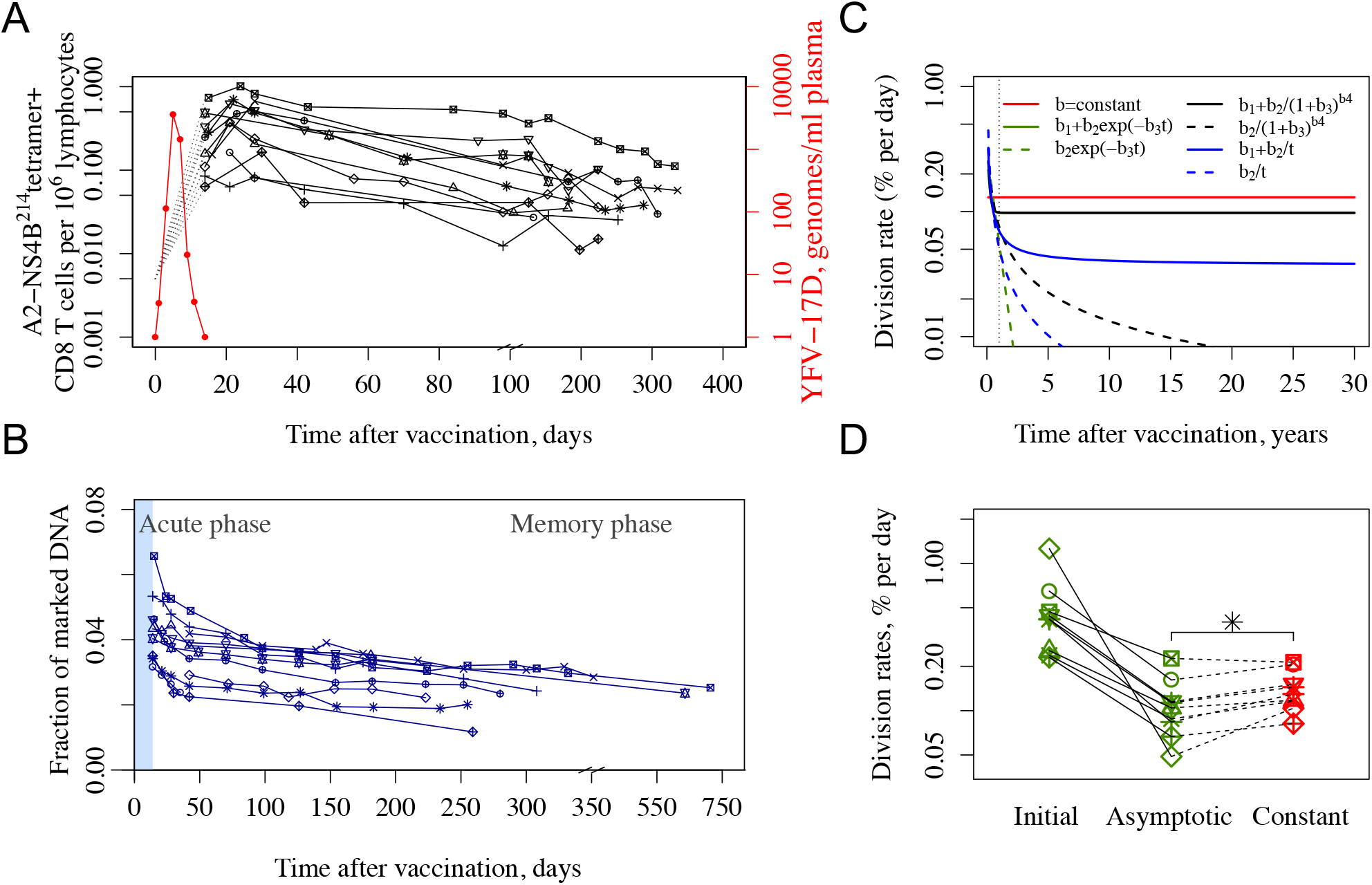
Dynamics and turnover of YFV-specific CD8 T cells in the first year after vaccination with the YFV-17D vaccine in Akondy study. Panel A shows the dynamics of the virus (red) and YFV-specific CD8 T cell numbers (black) in blood. Panel B shows the deuterium incorporation and die-away of label enrichment in YFV-specific CD8 T cells in individuals given D_2_O in their drinking water from day 0 to 14 post immunization (blue shaded region). Data in Panels A,B are from [23]. Panels C show the estimated division rate during 30 years for all models tested (different functions describing the change of division rate over time are shown and color-coded as red, black, green and blue, see also individual Donors fitting in S1 Fig). Panel D shows the division rate estimates with green symbols corresponding to initial and asymptotic division rates for the best model (*b*(*t*) = *b*_1_ + *b*_2_*exp*(−*b*_3_*t*)). The red symbols correspond to the division rates estimated from the constant division rate model. Each individual is shown by a different symbol. Solid and dotted lines connect the data for each individual obtained from the two models.

Akondy et al. [23] analyzed the dynamics and turnover of YFV-specific cells at memory phase (from day 42 post vaccination), assuming constant rates for the loss in the cell number and turnover of these cells. The estimated average rate of cell loss and rate of division of YFV-specific cells during the first year post vaccination was equal to 0.57 ± 0.08% per day (half-life of about 122 days) and 0.15 ± 0.09% per day (cells divide on average once every 462 days). Here, we consider a more refined analysis allowing the rate of loss of YFV-specific cells and its underlying cell division and death rates to change over time. We also focus on the data from day 42 post immunization to analyze the memory phase. This choice simplified the calculations, since by this time there was negligible D_2_O in the serum allowing us to neglect further deuterium incorporation into DNA of the cells.

We expect that division and death rates decline gradually and reach their asymptotic levels over the long-term, and thus we considered several different time-dependent functions to capture this. The fitting to the data was performed using non-linear mixed effect modeling framework implemented in Monolix Suite 2019 (Lixoft, France).

We first estimated the division rate, *b*(*t*), from the data for the fraction of marked DNA (Fig 1B). Four different functional forms of *b*(*t*) were used to fit the data, and results are shown in Fig 1C, D and S1 Fig. The model fits to the data were compared using Akaike Information Criterion (AIC) and Bayesian Information Criterion (BIC). The constant division rate model showed the worst fit. A general function *b*(*t*) = *b*_1_ + *b*_2_/(1 + *b*_3_*t*)^*b*_4_^ allows more flexibility with its parameter *b*_4_ influencing how fast *b*(*t*) approaches to asymptotic value *b*_1_. The fit to the data returned *b*_4_ = 26 with a major drop to a plateau level within a year after vaccination (solid black line in Fig 1C and S1B-D Fig). Interestingly, division rates estimated from the best fit model *b*(*t*) = *b*_1_ + *b*_2_*exp*(−*b*_3_*t*) and the general function model overlap at longer times (solid black and green lines in Fig 1C and S1C,D Fig). The asymptotic value *b*_1_ = 0.097 ± 0.053% per day (time between division of 715 days) of the best fit model is lower than 0.13 ± 0.044% per day (time between division of 533 days) estimated for the constant division rate model, and the difference is statistically significant (paired *t*-test *p*-value=0.006, Wilcoxon signed-rank test *p*-value=0.0078 for green and red symbols in Fig 1D). Functional form *b*(*t*) = *b*_1_ + *b*_2_/*t*, with one less parameter than the best fit model and ΔAIC=3, took longer to approach its asymptotic value of *b*_1_ = 0.0372% per day. Note that the data do not support models, in which cells stop dividing in the long-term (*b*_1_ = 0 in S1A Fig).

We next fitted both the data for the number of YFV-specific CD8 T cells and for the fraction of marked DNA (Fig 1A,B) simultaneously (see Methods). Similarly to the case of fitting the data for the fraction of marked DNA only discussed above, we first used the model with the more general form (division rate *b*(*t*) = *b*_1_ + *b*_2_/(1 + *b*_3_ * *t*)^*b*_4_^ and similar functional form for the death rate). In this general form, parameters *b*_4_ and *d*_4_ affect how quickly the corresponding rates approach their asymptotic values. Fitting it to the data returned interesting observations: relatively high value of parameter *b*_4_ for the division rate with relatively low corresponding parameter for the death rate (*d*_4_ = 0.043), consistent with the division rate approaching its asymptotic level much faster than the death rate. AIC for this general form model was high in comparison to other models tested due to extra parameters, and it was excluded from further analysis. The results for the different models tested are shown in Fig 2. We considered several simple functional forms for division and death rates where form *r*(*t*) = *r*_1_ + *r*_2_*exp*(−*r*_3_*t*) allowed faster decline to the asymptotic value and form *r*(*t*) = *r*_1_ + *r*_2_/*t* could account for a slower decline. In the simplest *exponential* model (Model 1, see Fig 2), the division and death rates are constants. Model 2 tests the hypothesis of two populations with constant division and death rates: a “stem cell like” population that may expand early during response but is relatively stable after day 42 with division and death rates equal to each other and another expanded population that is dying off with its *b* < *d*. Models 3-5 are different variations of the *progressive quiescence* model, where the overall population of YFV-specific cells progressively changes to establish long-term memory cells. Model 3 has both division and death rates change in the form of *r*(*t*) = *r*_1_ + *r*_2_*exp*(−*r*_3_*t*), Model 4 has both division and death rates in the form of *r*(*t*) = *r*_1_ + *r*_2_/*t*, and Model 5 has division rate as in Model 3 and death rate as in Model 4. For Models 2-5 we assume long-term division and death rates balancing each other as suggested by the mouse model [6, 7, 9], but also show how well the variants of Models 2-5 with unrestricted parameters (long-term division and death rates are not restricted) fit the data in parentheses in Fig 2A. Fig 2B shows how Models 1-5 capture both the change in the total number of cells and fraction of marked DNA in YFV-specific cells for each individual.

**Fig 2.**
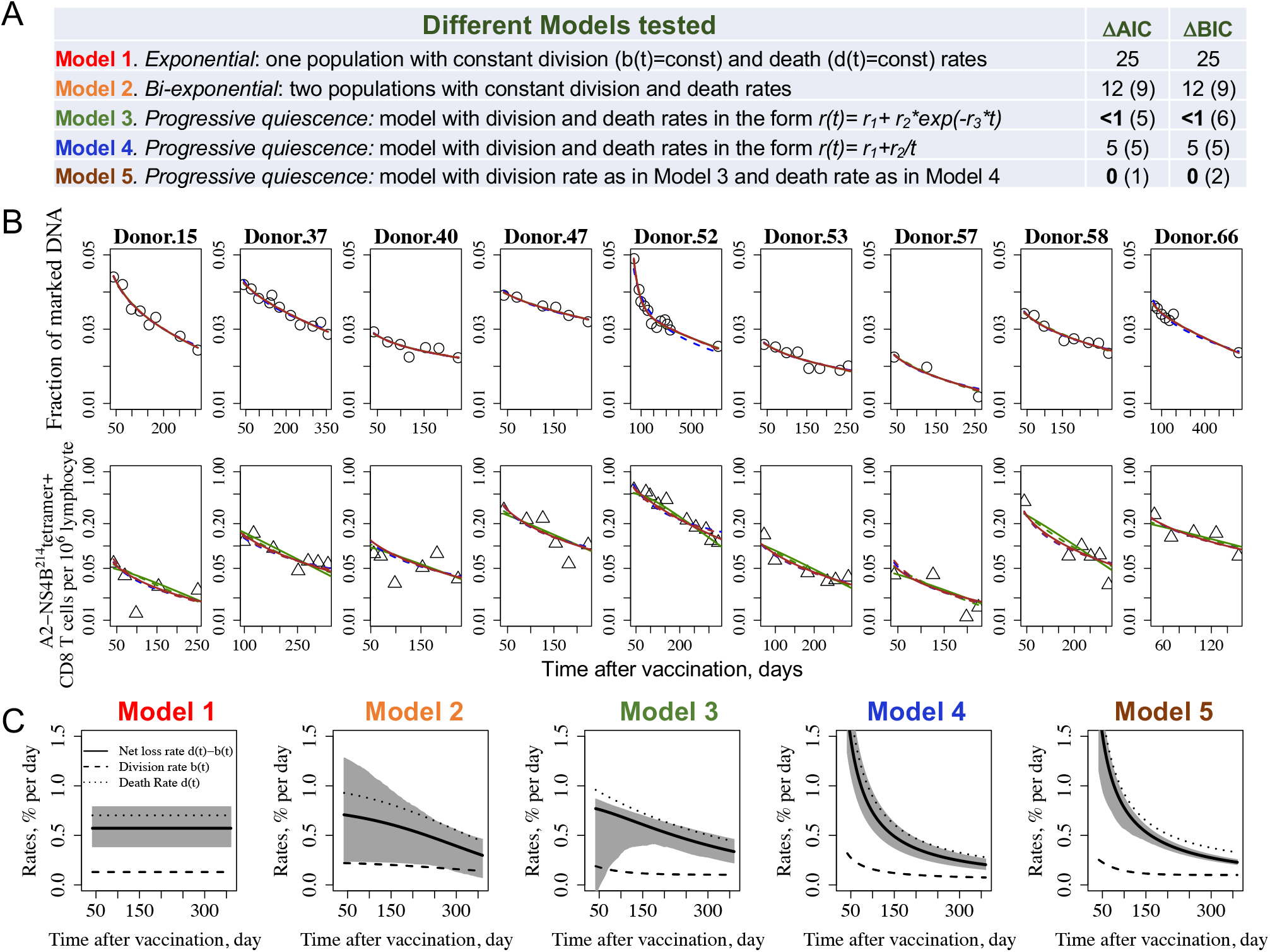
Changes in the tetramer positive YFV-specific CD8 T cells and their turnover over time. Panel A shows models describing different scenarios for changes in the division and death rates, b(t) and d(t), respectively, with time. In the simplest model (Model 1), the division and death rates are constant. Model 2 has two populations of cells with two different sets of constant division and death rates. In Models 3-5, the division and/or death rates can change from an initial value at day 42 to an asymptotic value long-term. Model 3 has both division and death rates as *r*(*t*) = *r*_1_ + *r*_2_*exp*(−*r*_3_*t*). Model 4 has both division and death rates modeled as *r*_1_ + *r*_2_/*t* and corresponds to decline in number of T cells described by power law, a functional form previously suggested to capture the waning of antibodies [28]. Model 5 has division rate as in Model 3 and death rate as in Model 4. We use a non-linear mixed effect modeling framework and AIC and BIC information criteria to examine how well these models fit the data from Akondy study [23, 29]. Model 2-5 have parameters restricted such that asymptotic division and death rates equal each other. ΔAIC and ΔBIC values are shown for these models in relation to best fit Model 5. ΔAIC and ΔBIC for unrestricted versions of these models are shown in parentheses. Panel B shows how Models 1 (red line), Model 2 (orange line), Model 3 (green line), Model 4 (blue line), and Model 5 (brown line) capture both the change in the total number of cells (data - triangles) and the deuterium labelling (fraction of marked DNA, data - circles) for each individual. Solid lines correspond to the models with long-term division and death rates equal each other and dashed lines correspond to the same models with unrestricted parameters (no requirement for long-term balancing in division and death rates). Panel C shows the predictions of models shown in Panel B for the division rate, death rate and overall net loss rate (i.e. death rate-division rate). The 95% CI for the net loss rate is shaded in grey (see Methods section).

As foreseeable, the data do not support the *exponential* decline model with constant division and death rates (ΔAIC=25 in comparison to best fit Model 5), nor does it support the *bi-exponential* decline model (Model 2) with two populations both having their division and death rates as constants (ΔAIC=12 for model with *b*_1_=*d*_1_ and ΔAIC=9 shown in parentheses for its unrestricted parameters version).

Fig 2C shows the estimates for how the division and death rates and the difference between them (which equals to the net loss rate) change with time for each model. While changes in division rates occur in a relatively narrow range in all models, changes in death rates could be significant and still allow for the fitting of the data reasonably well. Thus, analysis of the data from the Akondy study alone did not allow us to distinguish between the three models with ΔAIC≤5, but these models make different predictions for the long-term loss of YFV-specific CD8 cells, and next we will bring these predictions into contact with the data in Marraco study [26].

### Long-term maintenance of YFV-specific memory and model discrimination

The Marraco study [26] measured the numbers of YFV-specific T cells in individuals at different times during a period of 30 years following vaccination (Fig 3A). The black line shows the average decay rate for the entire dataset and gives a net loss of memory cells of 0.023% per day which corresponds to a “half-life” of about 8.25 years and consistent with previous estimates [11]. Interestingly, we find that the decay rate is not constant but rather declines with time. This can be seen by comparison of the red line which indicates that for the first 3 years the rate of loss of memory cells is much faster (0.18% per day, half life of only about 1 year) and green line showing that after 10 years the estimated net loss rate is much lower (0.009% per day). Fig 3B compares these estimates with Akondy study.

**Fig 3.**
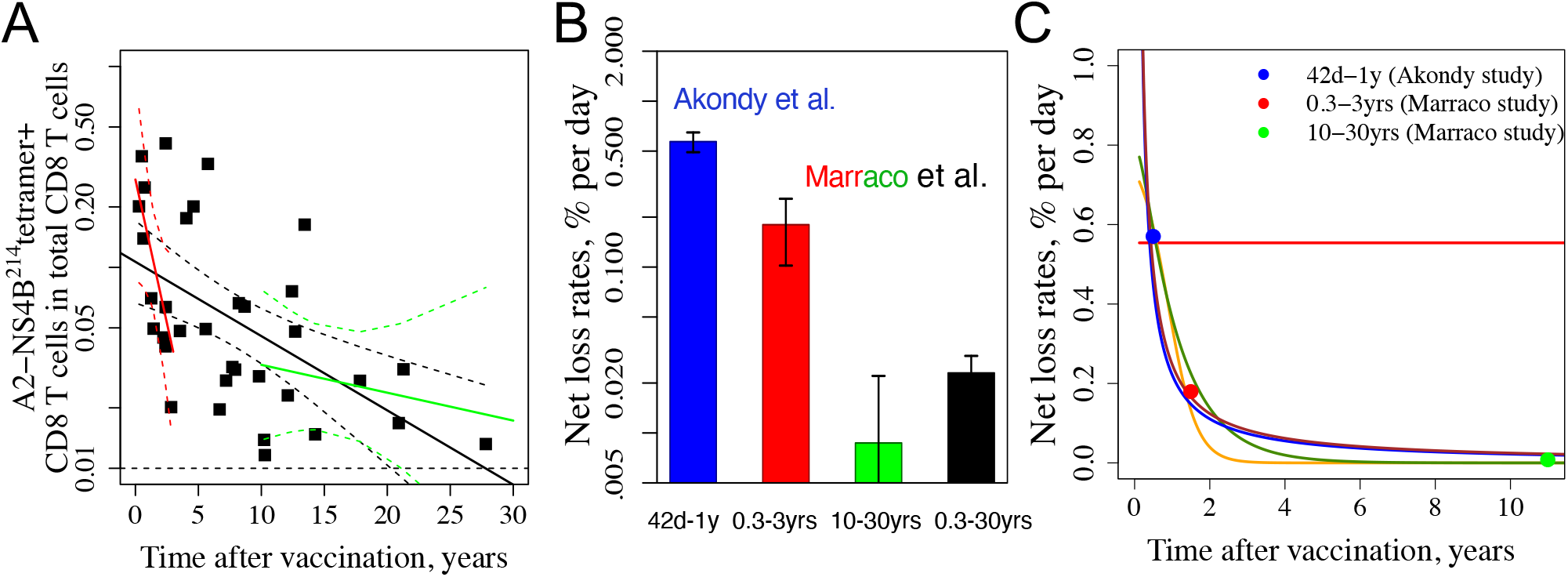
Dynamics of YFV-specific CD8 T cells over 30 years from the Fuertes-Marraco cross-sectional study [26]. Panel A shows the number of YFV-specific cells (on a log scale) as a function of time post vaccination. The slope gives the net rate of loss of the YFV-specific CD8 T cell response. The average rate of loss over the whole 30-years period is shown in the black line and the red and green lines show the initial (first 3 years) and long-term (years 10-30) rates of cell loss, respectively. Panel B summarizes the estimates from Panel A and compare them with Akondy study. Panel C shows the predictions from the Models 1-5 for the rate of cell loss, defined as the difference between the death and division rates at each time point, with Models 2-5 being more consistent with the experimental data. Filled circles are corresponding estimates from Panel B. Line colors: Model 1 - red, Model 2-orange, Model 3 - green, Model 4 - blue, Model 5 - brown.

We can bring predictions from our Models 1-5 into contact with the data in the cross-sectional Marraco study [26] in two different ways – by comparing (1) net loss rate of the YFV-specific cells and (2) the number of YFV-specific cells with time. Fig 3C shows predictions for all models for the net loss rate (colored lines) together with estimates from the two experimental studies (circles, colors correspond to the bars in Fig 3B). Four models shown in orange (Model 2), green (Model 3), blue (Model 4) and brown (Model 5) lines are consistent with the data. The predicted rate of loss of the YFV-specific cell population in these four models gradually falls over time, and a relatively stable memory cell population with a very low net loss rate is reached several years after immunization. The change in the rate of net loss of CD8 T cells is due to a slow decline of the death rate of the population over this time frame and only a modest, less than two fold, change in the division rate (see Fig 2C).

Predictions of the models for a decline in the number of YFV-specific CD8 T cells together with the data from Akondy (open squares) and Marraco (filled squares) studies are shown in Fig 4. Residual Sums of Squares (RSSs) indicate how well the predictions fit the Marraco data. Model 4 and Model 5 fit data the best followed by Model 3 and then by Model 2. Model 2 has two populations both having constant division and death rates, and the division and death rates are balancing each other long-term for one of the populations. Model 2 shows a poor fit to the Akondy’s data (ΔAIC=12) and RSS value from its prediction of Marraco data do not support it as well. In Model 4, both division and death rates are in the form of *r*(*t*) = *r*_1_ + *r*_2_/*t*, while Model 5 uses *b*(*t*) = *b*_1_ + *b*_2_*exp*(−*b*_3_*t*) for the division rate and *d*(*t*) = *d*_1_ + *d*_2_/*t* for the death rate. While both Models 4 and 5 fit the Marraco’s data well and their RSS values are not significantly different (paired t-test on their squared residuals has p-value = 0.87), they make different predictions regarding the long-term division rate. Model 4 predicts 0.043% per day (very slow doubling time ≈ 1600 days) while Model 5 predicts 0.1% per day (turnover rate ≈ 700 days). Three factors suggest that Model 5 may better describe the data overall: (1) better fit of deuterium incorporation data alone (S1 Fig), (2) better fit of the data in Akondy study (Fig 2A), and (3) it is more consistent with the independent estimate of division rate as 0.15 ± 0.045% per day using data on D_2_O intake at 4-9 months post vaccination [23] (S3 Fig).

**Fig 4.**
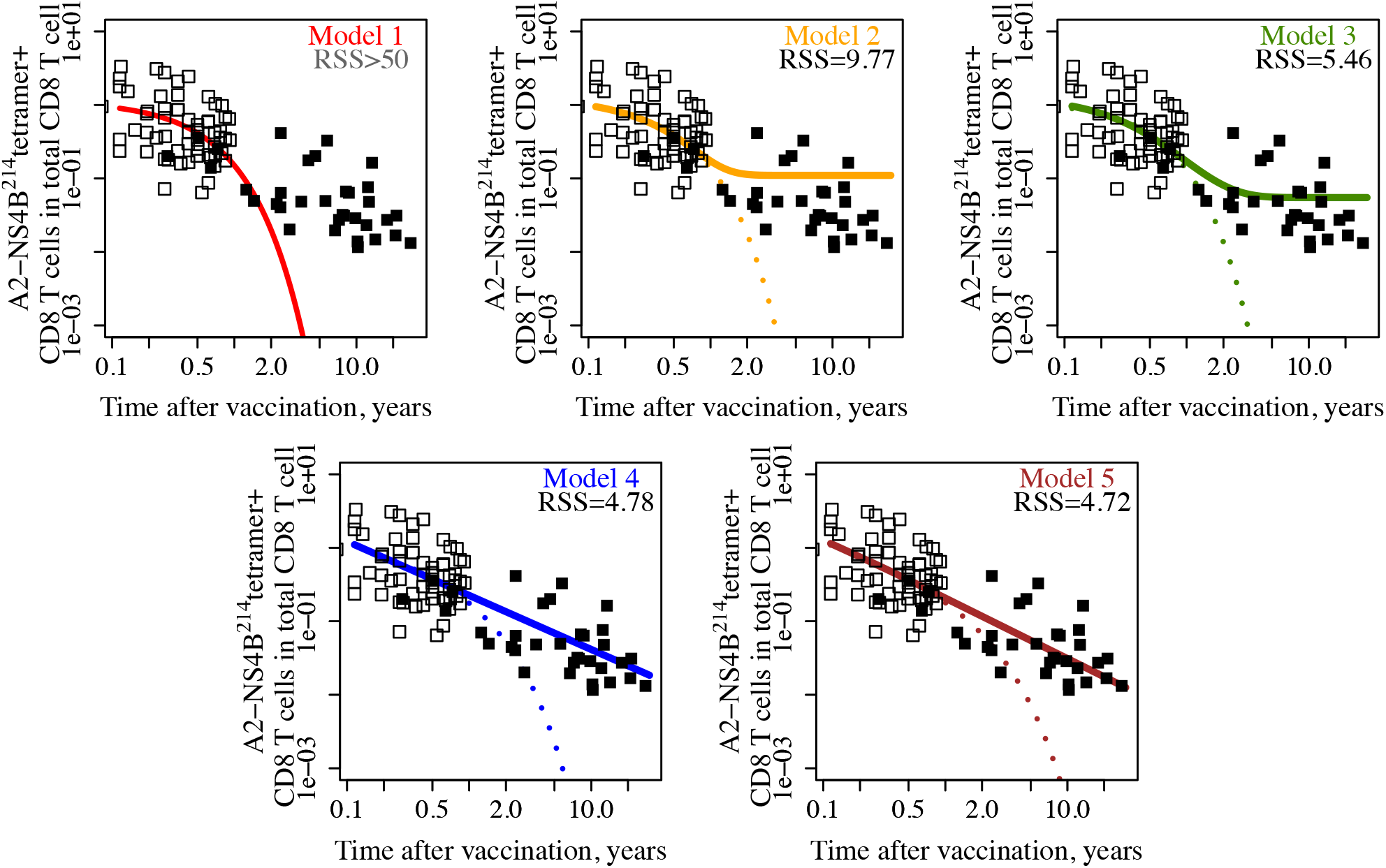
Integrated analysis of the data suggests that Models 4 and 5 predict the data in Marraco study better than the other models. Data from the Akondy study [23] and the Marraco study [26] are shown as open and closed squares, respectively. For each model, we use fit to the Akondy data to predict the number of YFV-specific cells up to 30 years after vaccination and compare it with Marraco data using Residual Sums of Squares (RSSs). Dotted lines in panels with Model 2-5 show predictions from the corresponding models with unrestricted parameters. RSSs for these models are more than 50 (not shown) indicating a poor fit. As YFV-specific cells in the Akondy study were measured as % in PBMCs and in the Marraco study as % in CD8 T cells, we converted the cell numbers from fitting Akondy data and model predictions using the following reasonings. Since 45-70 % of PBMCs are T cells and about 30 % of them are CD8 T cells, we used a conversion factor equal to 0.575o0.3≈ 0.17. Plotting how this result depends on the conversion factor (S2 Fig) shows that Models 4 and 5 consistently show a better fit. Models colors are as in Fig 2: Model 1 - red, Model 2 - orange, Model 3 - green, Model 4 - blue, Model 5 - brown.

### Power law predicts change in YFV-specific cells

Changes in the net loss rate over time define the number of YFV-specific cells. Interestingly, the number of cells in our two best models for prediction of the YFV-specific cells are described by power law functions *exactly* in Model 4 (division rate *b*(*t*) = 0.00043 + 0.12/*t*, death rate *d*(*t*) = 0.00043 + 0.85/*t*, so *d*(*t*) − *b*(*t*) = 0.73/*t* and frequency of YFV-specific cells *N*(*t*) ∝ *t*^−0.73^ for t≥42 days) or *approximately* for Model 5 (*b*(*t*) = 0.001 + 0.0038 * *exp*(−0.022*t*), *d*(*t*) = 0.001 + 0.82/*t*, so *d*(*t*) − *b*(*t*) ≈ 0.82/*t* as division rate *b*(*t*) ≈ 0.001 after first year and asymptotic division and death rates are equal each other, so *N*(*t*) ≈ *c* · *t*^−0.82^ where *c* is a constant). We found that fitting longitudinal data in Fig 1A with a *simple power law* model from day 42 using mixed-effect framework can predict the number of YFV-specific cells for the decades after immunization. Note that we are using only YFV-specific T cell data and no DNA enrichment data (*dN*/*dt* = −*k*/*t* · *N*, *N*(*t*) ∝ *t*^−*k*^, *k*=0.76). Fig 5 shows the simple power law model’s predictions (purple line) for Marraco data in comparison to Models 4 (blue line) and Model 5 (brown line) predictions.

**Fig 5.**
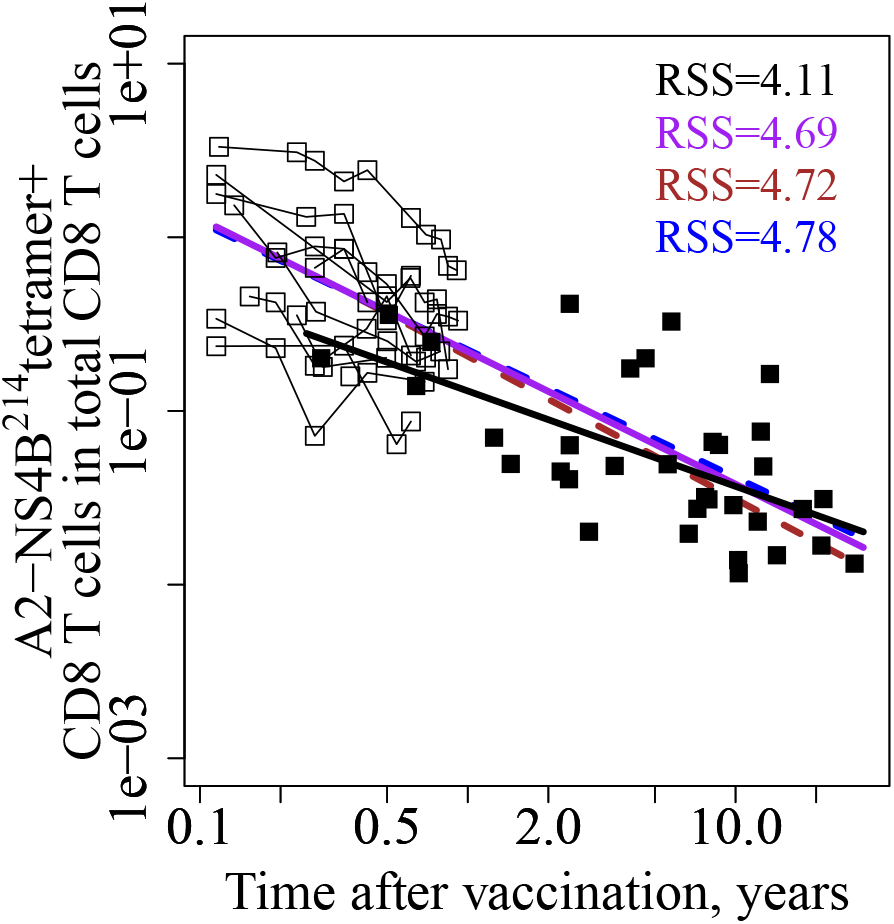
Predictions for Marraco’s data from Model 4 (blue line), Model 5 (brown line), and simple power law model (purple line). Data from the Akondy study [1] and the Marraco study [2] are shown as open and closed squares, respectively. Data points for each individual donor in Akondy study are connected by thin black line. For Models 4 and 5, we fit the models to the Akondy data (Fig 1) from day 42 to 1 year using mixed-effect framework to estimate the population parameters and based on these population parameters predict the number of YFV-specific cells up to 30 years after vaccination. We compare models predictions with Marraco data using RSSs (RSS value colors matched the corresponding lines). Unlike Models 4 and 5, simple power law model is using only YFV-specific T cells data (Fig 1A) and no DNA enrichment data. Use of mixed-effect framework is essential for accurate prediction for all three models. Additionally, power law fit for Marraco’s data only is shown with a black line.

## Discussion

We developed a set of mathematical models to explore how the rates of division and death of YFV-specific CD8 T cells change with time after immunization with YFV-17D that causes a mild acute infection. Analysis of the Akondy data alone allowed us to reject the commonly used exponential (ΔAIC=25) and biexponential (ΔAIC=12) decline models (Fig 2). Furthermore, there was poor support for models where the division rate asymptotes to zero in the long-term (S1 Fig) suggesting that long-term memory in humans is maintained by a population undergoing turnover. This result is qualitatively similar to memory cells in mice [1, 6, 9], albeit the turnover of memory cells in mice (‘doubling time’ ≈50 days) is much faster than in humans (our estimated ‘doubling time’ ≈700 days). Combined analysis of the Akondy and Marraco data allowed us to discriminate between different models. Model 5, where division rate is in form *b*_1_ + *b*_2_*exp*(−*b*_3_*t*) and death rate is in form *d*(*t*) = *d*_1_ + *d*_2_/*t* and they asymptote to the same level (*b*_1_ = *d*_1_), albeit at different rates, not only fits best the Akondy data but the Marraco data as well. Models in which the rate of cell loss follows a power law (exactly or approximately) are able to accurately predict the frequencies of YFV-specific T cells up to 30 years post vaccination.

An earlier study by Teunis et al. [28] suggested that the waning of antibodies to pertussis can be described better by a power law in comparison with single or bi-exponential forms of decay that were used earlier [30–32]. Our models suggest that the power law can also be used to describe the waning of CD8 T cell immunity. The gradual decline in the decay rate of CD8 T cell immunity is in line with previous studies for CD8 T cell memory following vaccinia immunization [10, 11].

There have been a number of earlier studies using deuterium labelling to quantify the turnover of T cells in humans [17, 18, 20, 21, 33–36]. Most of these studies identified CD8 memory cells on the basis of the CD45RO^+^ marker [17, 33, 36] and suggested that cells of this phenotype had a proliferation rate in the range of 0.6% to 2% per day [18, 33]. Further analysis suggested the disparity in these estimates could be reduced by using biphasic [34] or multi-exponential models [18], which suggests that there are subpopulations of CD45RO^+^ cells with different rates of turnover. Hellerstein et al. [34] analyzed bulk CD8 T-cell turnover in healthy subjects during longer-term labeling (between weeks 5 – 9) with D_2_O or longer term die-away (between weeks 3-7) after pulse labeling with 2H-glucose and reported 2% replacement per week by both approaches, or a division rate of 0.3% per day. More detailed phenotypic analysis of memory cells showed not only that there are different subpopulations of CD45RO^+^ memory cells, but also that at later times following immunization there are CD45RO^-^ memory cells which share many surface markers (CD45RO^-^, CCR7^+^, CD28^+^, CD127^+^) with naive CD8 T cells, but can be distinguished from naive cells on the basic of CXCR3, CD31, CD11a and CD95 [23, 37]. These cells have been termed stem-cell like memory cells or T_SCM_, and it has been shown that the fraction of cells with this phenotype increases with time following YFV immunization [23]. Consequently, the studies focusing on CD45RO^+^ cells will not describe the turnover of antigen-specific long-term memory CD8 T cells and they caution against attempts to identify a memory cell population with just a few surface markers.

Our estimate for the long-term division rate of YFV-specific cells is 0.1% per day and could be attributed to T_SCM_ population, as majority of cells detected decades after vaccination have T_SCM_ phenotype [23]. It is lower than previous estimate of 0.6-7% per day [21] and positions the division rate for T_SCM_ cells between the rates for naive (0.03-0.06% per day [17, 21]) and memory (0.3-2% per day [18, 33, 34, 38]) cells.

There are caveats and limitations associated with our analysis, and we will now briefly discuss the important ones. As with many previous studies, we focus on data obtained from cells circulating in the blood and assume that the changes in the populations in the blood are representative of changes elsewhere. This is likely to be a reasonable assumption based on studies that demonstrate recirculation of memory T cells, but our analysis will not apply to resident memory populations that can play an important role in some infections [39]. A second limitation is that memory is associated with changes in both the number of antigen-specific CD8 T cells and a change in their phenotype, and this study focuses exclusively on the former. Recent studies have shown that CD8 T cells remain poised after primary response to have more rapid responses to repeat infections [8, 23], and the contribution to protection of phenotypic changes versus changes in numbers of antigen-specific cells has yet to be quantitatively evaluated. Finally, our studies consider turnover of the YFV-specific CD8 T cells as a whole – and at this stage lack of data precludes analysis of the differentiation pathways between different subpopulations which are likely to have different division and death rates. Deuterium labelling studies that follow cells of different phenotypes with a defined antigenic specificity will be needed to address this question in humans.

The immune system maintains immune memory and also responds to antigenically novel infections. Our results suggest that CD8 T cell memory declines approximately in accordance with a power law. In the case of the yellow fever vaccine, exponential decline at the rate observed 6 weeks to 1 year post vaccination (0.55% per day) would lead to dramatically fewer memory cells long term (>6 fold lower after only 10 years) compared to what is actually observed. Hence this exponential decline model is not only empirically inaccurate but potentially incompatible with longlasting protection via immune memory. In contrast, a power law is consistent with both the initial decline and long term maintenance of memory CD8 T cells. Notably, a power law fit to the first year of data accurately predicted memory CD8 T cell numbers over three decades.

For new vaccines and emerging infections, such as SARS-CoV-2, where long term immune memory data is not yet available, power laws may be a useful alternative to exponential and biexponential models for predicting future waning of the immune response.

## Methods

We build our model on previously published studies [17, 40]. The model equations are:

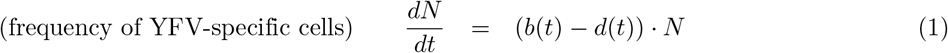

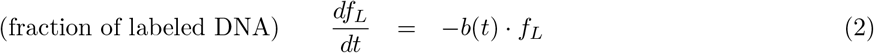

We assume that both division rate *b*(*t*) and death rate *d*(*t*) change with time and asymptote to non-zero values, unless indicated otherwise. The asymptotic value of the division rate was used to calculate doubling time as *ln*2/*b*. Functions used to model division and death rates are shown in Fig 2A and S1A Fig. Since we focus on fitting the data from day 42 post immunization (28 days after administration of deuterium labelled water was stopped), the heavy water was largerly washed out (half-time approximately 9 days [23]) allowing us to neglect further deuterium incorporation into DNA of the cells and to simplify the analysis.

For the two population bi-exponential decline model, we used two sets of equations each similar to the one population exponential decline model and fit the data shown in Fig 1A,B with *N* = *N*_1_ + *N*_2_ and *f_L_* = (*f*_*L*1_ · *N*_1_ + *f*_*L*2_ · *N*_2_)/(*N*_1_ + *N*_2_), respectively, where subscripts correspond to these two populations.

Non-linear mixed effect models were implemented using Monolix Suite 2019 (Lixoft, France). Both frequency of YFV-specific CD8 T cells and fraction of labeled DNA data were fit simultaneously for all donors. We assumed that model parameters are log-normally distributed. Fixed and random effects values were estimated for the model parameters in the functions for the division and death rates and initial condition values for model variables. The estimation of the population parameters was performed using the Stochastic Approximation Expectation-Maximization (SAEM) algorithm. Model comparison was based on information criteria such as the Akaike information criterion (AIC) [41] and Bayesian information criterion (BIC) [42]. To add 95% CIs in Fig 2C and S3B Fig, we ran 1000 simulations for each model with Simulx (Lixoft, France) using the corresponding population parameters obtained from fitting the data in Monolix.

To plot the data in Fig 3A we digitized the data from Figure 1B in the Fuertes-Marraco cross-sectional study [26] using WebPlotDigitizer (https://apps.automeris.io/wpd/).

## Acknowledgments

This work was supported by National Institutes of Health U01 HL139483 and U01 AI150747.

