## Supplemental figures for "Dynamics and turnover of memory CD8 T cell responses following yellow fever vaccination"

### Supporting information

**S1 Fig. Estimation of division rate of YFV-specific cells from the four models used to fit the fraction of marked DNA shown in Fig 1B.** Panel A shows the Table with different functions describing the change of division rate over time in different models color-coded as red, black, green and blue. Corresponding ΔAIC and ΔBIC are listed in comparison to the best model $b(t) = b_1 + b_2 exp(-b_3 t)$. Cases where long-term division rate is equal to zero ($b_1$=0) showed worse fit for all functional forms as shown by ΔAIC and ΔBIC in the brackets with grey color for each model. Panel B shows the fitting results for individual Donors using data from day 42 upward for each model with colors as in Panel A. Panels C and D show the estimated division rate during 1 year and long-term during 30 years for all models tested. Vertical dotted line in Panel D corresponds to 1 year.

A

| Different Models tested | ΔAIC | ΔBIC |
|---|---|---|
| Constant division rate, b(t)=constant, solid red line | 27 | 27 |
| General form $b(t)= b_1+b_2/(1+b_3*t)$^$b_4$, solid black line (dashed black line for $b_1$=0) | 5 (15) | 5 (15) |
| $b(t)=b_1+b_2*exp(-b_3*t)$, solid green line (dashed green line for $b_1$=0) | 0 (5) | 0 (4) |
| $b(t)= b_1+b_2/t$, solid blue line (dashed blue line for $b_1$=0) | 3 (4) | 3 (3) |

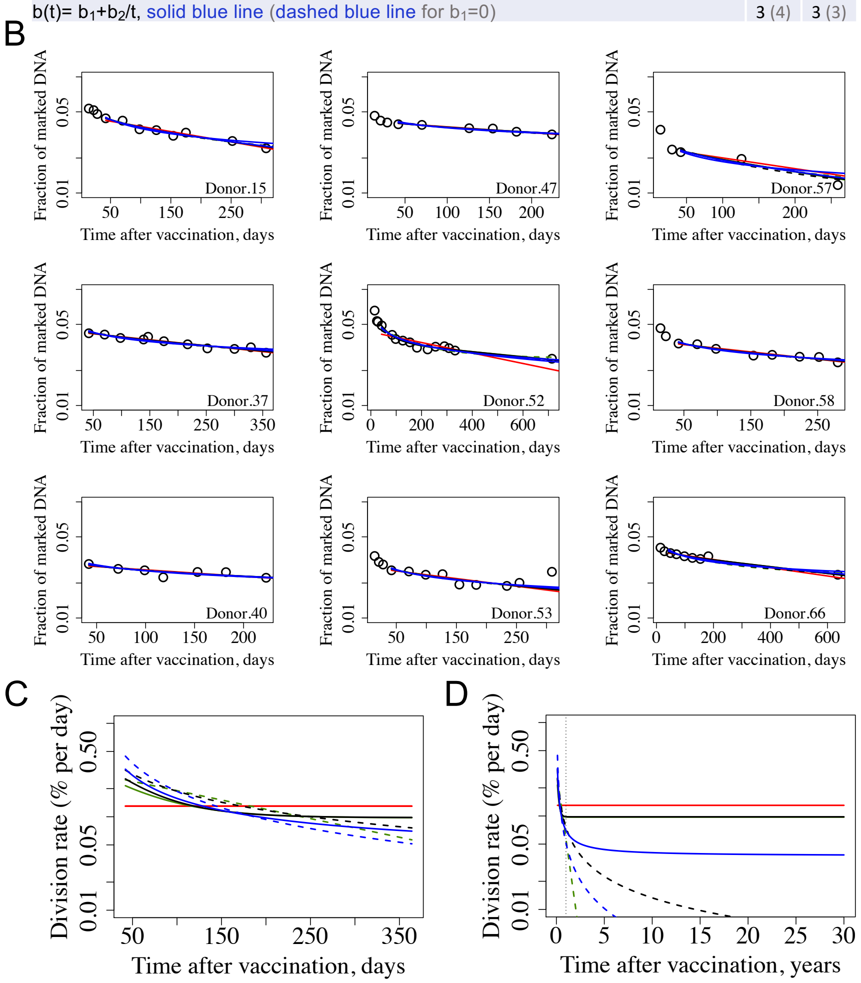

**S2 Fig. Dependence of Sums of Squares (RSS) for Models 2-5 on conversion factor.** YFV-specific cells in the Akondy study were measured as % in PBMCs and in the Marraco study as % in CD8 T cells, thus we converted the cell numbers from fitting Akondy data and model predictions using the following reasonings. Since 45-70 % of PBMCs are T cells and about 30 % of them are CD8 T cells, we used a conversion factor equal to 0.575*0.3 ≈ 0.17 in Fig 4. Plotting how this result depends on the conversion factor in a range of between 0.135 and 0.21 shows that Models 4 and 5 consistently show a better fit. Models colors are as in Fig 2: Model 2 - orange, Model 3 - green, Model 4 - blue, Model 5 - brown.

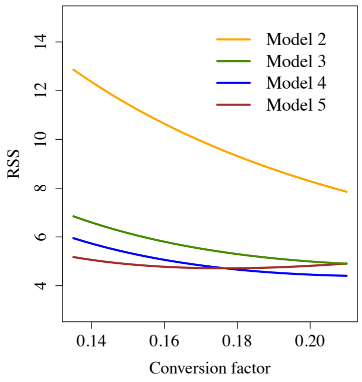

**S3 Fig. Predictions from Models 4 and 5 compared to estimate for deuterium incorporation at late memory stage.** Panel A shows deuterium incorporation in six Donors with heavy water consumption for 8 weeks with start at 4-9 months after YFV immunization (Study 3 in [23]). Data for different Individuals are shown by different symbols. Pink shaded area indicates the period of heavy water consumption for each Donor. Panel B shows how corresponding division rates (with their 95% confidence intervals (CI)) change with time. Asymptotic division rate for Model 4 was estimated as 0.043% per day (95% CI 0.012-0.148) and for Model 5 as 0.1% per day (95% CI 0.041-0.235). Akondy study analyzed the deuterium incorporation at the memory stage shown in Panel A [23], and the corresponding estimate for division rate equal to 0.15±0.045% per day is shown by rectangle with 4-19 month length on x-axis and standard deviation estimation on y-axis.

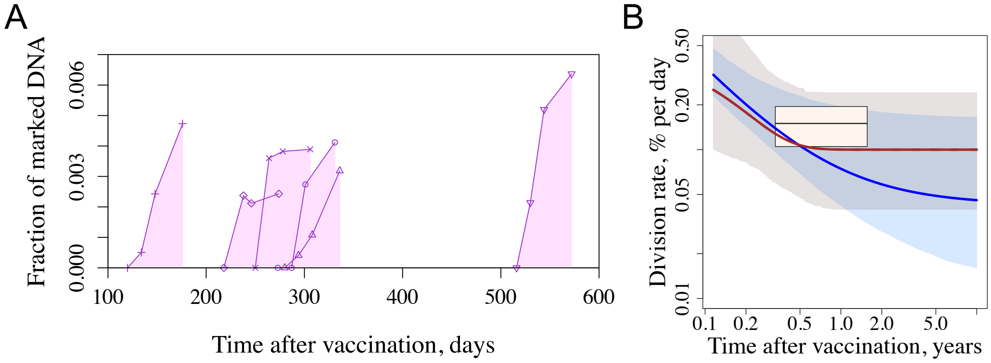
